# Convergent origin and accelerated evolution of vesicle-associated RhoGAP proteins in two unrelated parasitoid wasps

**DOI:** 10.1101/2023.06.05.543686

**Authors:** Dominique Colinet, Fanny Cavigliasso, Matthieu Leobold, Appoline Pichon, Serge Urbach, Dominique Cazes, Marine Poullet, Maya Belghazi, Anne-Nathalie Volkoff, Jean-Michel Drezen, Jean-Luc Gatti, Marylène Poirié

## Abstract

Animal venoms and other protein-based secretions that perform a variety of functions, from predation to defense, are highly complex cocktails of bioactive compounds. Gene duplication, accompanied by modification of the expression and/or function of one of the duplicates under the action of positive selection, followed by further duplication to produce multigene families of toxins is a well-documented process in venomous animals. This evolutionary model has been less described in parasitoid wasps, which use maternal fluids, including venom, to protect their eggs from encapsulation by the host immune system. Here, we evidence the convergent recruitment and accelerated evolution of two multigene families of RhoGAPs presumably involved in virulence in two unrelated parasitoid wasp species, *Leptopilina boulardi* (Figitidae) and *Venturia canescens* (Icheumonidae). In both species, these RhoGAPs are associated with vesicles that act as transport systems to deliver virulence factors, but are produced in different tissues: the venom gland in *Leptopilina* sp. and the ovarian calyx in *V. canescens*. We show that the gene encoding the cellular RacGAP1 is at the origin of the virulent RhoGAP families found in *Leptopilina* sp. and *V. canescens*. We also show that both RhoGAP families have undergone evolution under positive selection and that almost all of these RhoGAPs lost their GAP activity and GTPase binding ability due to substitutions in key amino acids. These results suggest an accelerated evolution and functional diversification of these vesicle-associated RhoGAPs in the two phylogenetically distant parasitoid species. The potential new function(s) and the exact mechanism of action of these proteins in host cells remain to be elucidated.

## Introduction

Gene duplication is recognized as an important evolutionary process because it drives functional novelty (Chen et al., 2013; Long et al., 2013). The importance of gene duplication for genetic and functional innovation was popularized by Ohno, who postulated that one of the two duplicate copies could evolve a new function through mutation, while the other would be responsible for the ancestral function (Ohno 1970). Since then, there have been numerous examples of duplication followed by neofunctionalization of one of the two paralogs with critical roles in fundamental biological processes (Kaessmann 2010; Magadum et al. 2013, Copley 2020). In particular, the formation of multigene families by repeated gene duplication is a widely studied evolutionary process in several groups of venomous animals (Fry et al., 2009; Wong & Belov 2012; Casewell et al., 2013), but also in some parasites (Akhter et al., 2012; Arisue et al., 2020). Such repeated duplication events of venomous or virulence protein-coding genes are often accompanied by significant copy divergence through positive selection, allowing the acquisition of new functions. In this work, we studied the process of accelerated evolution by duplication and divergence of two multigene families encoding RhoGAPs presumably involved in virulence in two phylogenetically distant parasitoid wasp species, *Leptopilina* species of the family Figitidae and *Venturia canescens* of the family Icheumonidae.

The development of parasitoid wasps occurs at the expense of another arthropod, whose tissues are consumed by the parasitoid larvae, usually resulting in the host death (Godfray 1994). For endoparasitoids that lay eggs inside the host body, the host immune defense against parasitoids is usually based on the formation of a multicellular melanized capsule around the parasitoid egg, resulting in its death (Carton et al., 2008). To escape encapsulation, parasitoids have evolved several strategies, the main one being the injection of maternal fluids together with the egg into the host at the time of oviposition (Pennacchio & Strand 2006; Poirié et al., 2009). These maternal fluids contain (i) proteins synthesized in the venom glands (Asgari & Rivers 2010; Poirié et al., 2014), some of which, as in *Leptopilina,* can be associated with extracellular vesicles called venosomes and allow their transport to the targeted immune cells (Gatti et al., 2012; Wan et al., 2019), (ii) proteins synthesized in the ovarian calyx cells and associated with particles of viral origin devoid of nucleic acid and named virus-like particles (VLPs) as in *V. canescens* (Reineke et al., 2006; Gatti et al., 2012; Pichon et al., 2015), and (iii) polydnaviruses (PDVs) integrated into the parasitoid genome and found in some groups of parasitoids belonging to the families Braconidae and Ichneumonidae (Drezen et al., 2014). Like VLPs, PDVs are synthesized in the ovarian calyx cells and are present in the ovarian fluid along with the egg. These particles are unique in that they are formed by the integrated viral machinery, but carry circular double-stranded DNA molecules that contain virulence genes that will be expressed in the host cells (Drezen et al., 2014). The process of accelerated evolution by duplication and divergence has been described for some of these virulence genes carried by PDVs, suggesting functional diversification of the proteins produced (Serbielle et al., 2008; Serbielle et al., 2012; Jancek et al., 2013).

*Leptopilina boulardi* is a parasitoid wasp of *Drosophila* for which two lines have been well characterized (Dubuffet et al., 2009). The ISm line of *L. boulardi* is highly virulent against *Drosophila melanogaster* but is unable to develop in *Drosophila yakuba* whereas the ISy line can develop in both *Drosophila* species but its success depends on the resistance/susceptibility of the host (Dubuffet et al., 2009). One of the most abundant proteins in the venom of the *L. boulardi* ISm line is LbmGAP (previously named LbGAP), which belongs to the Rho GTPase activating protein (RhoGAP) family (Labrosse et al., 2005; Colinet et al., 2013). A LbmGAP ortholog named LbyGAP (previously named LbGAPy) is also present in the venom of the *L. boulardi* ISy line, but in lower amounts (Colinet et al., 2010; Colinet et al., 2013). While RhoGAPs are usually intracellular proteins composed of several different domains (Tcherkezian & Lamarche-Vane 2007), LbmGAP and LbyGAP contain only one RhoGAP domain preceded by a signal peptide that allows its secretion. LbmGAP has been shown to be associated with and transported by venosomes to target host hemocytes (Wan et al., 2019). LbmGAP and LbyGAP specifically interact with and inactivate two *Drosophila* Rho GTPases, Rac1 and Rac2 (Colinet et al., 2007; Colinet et al., 2010), which are required for hemocyte proliferation in response to parasitism, hemocyte adhesion around the parasitoid egg, or the formation of intercellular junctions necessary for the capsule formation (Williams et al., 2005; Williams et al., 2006). Combined transcriptomic and proteomic analyses have identified eight additional RhoGAP domain-containing venom proteins in addition to LbmGAP and LbyGAP for both *L. boulardi* lines, suggesting that a multigene family has derived from repeated duplication (Colinet et al., 2013). One of these venom RhoGAPs, LbmGAP2 (previously named LbGAP2), has also been shown to be associated with venosomes in *L. boulardi* ISm and to be released with LbmGAP in host hemocytes (Wan et al., 2019). Interestingly, three RhoGAP domain-containing venom proteins have also been identified in the closely related species *Leptopilina heterotoma* (Colinet et al., 2013), suggesting that the recruitment of RhoGAPs in the venom arsenal may have occurred before the separation of the two species.

*Leptopilina* venom RhoGAPs are not the only example of the possible use of a protein from this family in parasitoid virulence (Reineke et al., 2006; Du et al., 2020). In the Ichneumonidae *V. canescens*, a RhoGAP domain-containing protein known as VLP2 (named VcVLP2 in this paper) was found to be associated with VLPs formed in the nucleus of ovarian calyx cells (Reineke et al., 2006; Pichon et al., 2015). VLPs, which package proteic virulence factors wrapped into viral envelopes, are then released into the ovarian lumen to associate with eggs and protect them from the host immune response by as yet unknown mechanisms (Feddersen et al., 1986). Species of the genus *Leptopilina* (superfamily Cynipoidea) as well as *V. canescens* (superfamily Ichneumonoidea) belong to the group of parasitoid wasps, which has been shown to be monophyletic (Peters et al., 2017). They are distantly related, with their last common ancestor dating back to the early radiation of parasitoid wasps, more than 200 million years ago (Peters et al., 2017). The presence of RhoGAP family proteins in the maternal fluids of parasitoid wasps has not been described outside of *Leptopilina* species and *V. canescens*. Overall, these observations suggest convergent recruitment of proteins belonging to the same family and injected into the host with the egg in two phylogenetically distant parasitoid species.

In this work, we showed that VcVLP2 in *V. canescens*, like LbmGAP in *Leptopilina*, is not unique but is part of a multigene family of virulent RhoGAPs associated with extracellular vesicles. RhoGAPs from different organisms are grouped into distinct subfamilies based on similarities in RhoGAP domain sequence and overall multi-domain organization (Tcherkezian & Lamarche-Vane 2007). Our analyses indicate that an independent duplication of the RacGAP1 gene, a member of the large RhoGAP family, was the origin of the two virulent RhoGAP multigene families, one found in *Leptopilina* and the other in *V. canescens*. We then performed comparative analyses to understand the duplication events at the origin of these two virulent RhoGAP multigene families. Finally, we demonstrated evolution under positive selection for both virulent RhoGAP multigene families in *L. boulardi* and *V. canescens,* suggesting accelerated evolution and functional diversification.

## Methods

### Biological material

The origin of the *L. boulardi* isofemale lines ISm (Lbm; Gif stock number 431) and ISy (Lby; Gif stock number 486) has been described previously (Dupas et al., 1998). Briefly, the ISy and ISm founding females were collected in Brazzaville (Congo) and Nasrallah (Tunisia), respectively. The *L. heterotoma* strain (Lh; GIF stock number 548) was collected in southern France (Gotheron). The Japanese strain of *Leptopilina victoriae* (Lv), described in Novković et al., 2011, was provided by Pr. M. T. Kimura (Hokkaido University, Japan). All parasitoid lines were reared on a susceptible *D. melanogaster* Nasrallah strain (Gif stock number 1333) at 25 °C. After emergence, the wasps were maintained at 20 °C on agar medium supplemented with honey. All experiments were performed on 5-to 10-day-old parasitoid females.

### Phylogeny of members of the Cynipoidea superfamily

The phylogeny of selected members of the Cynipoidea superfamily was constructed using internal transcribed spacer 2 (ITS2) sequences available at NCBI (Supplementary Table S1). Multiple sequence alignment was performed using MAFFT with the--auto option (Katoh & Standley 2013). Poorly aligned regions were removed using trimal with the-automated1 option (Capella-Gutierrez et al., 2009). Phylogenetic analysis was performed using maximum likelihood (ML) with IQ-TREE (Minh et al., 2020). The alignments obtained before and after using Trimal are shown in Supplementary Dataset 1.

### Search for candidate venom RhoGAPs in Leptopilina transcriptomes

*L. victoriae* venom apparatus, corresponding to venom glands and associated reservoirs, were dissected in Ringer’s saline (KCl 182 mM; NaCl 46 mM; CaCl_2_ 3 mM; Tris-HCl 10 mM). Total RNA was extracted from 100 venom apparatus using TRIzol reagent (Invitrogen) followed by RNeasy Plus Micro Kit (QIAGEN) according to the manufacturers’ instructions. The quality of total RNA was checked using an Agilent BioAnalyzer. Illumina RNASeq sequencing (HiSeq 2000, 2 × 75 pb) and trimming were performed by Beckman Coulter Genomics. The raw data are available at NCBI under the BioProject ID PRJNA974978. Sequence assembly was performed using the “RNASeq de novo assembly and abundance estimation with Trinity and cdhit” workflow available on Galaxy at the BIPAA platform (https://bipaa.genouest.org).

*Leptopilina clavipes* (whole body), *Ganaspis* sp. G1 (female abdomen), *Andricus quercuscalicis* (whole body) and *Synergus umbraculus* (whole body) transcriptomes were obtained from NCBI under accession numbers GAXY00000000, GAIW00000000, GBNY00000000 and GBWA00000000, respectively. Coding regions in the transcripts were identified and translated using TransDecoder (Haas et al., 2013), which is available on the BIPAA Galaxy platform. Searches for RhoGAP domain-containing sequences in translated coding sequences were performed using hmmsearch from the HMMER package (Eddy 2009) with the RhoGAP (PF00620) HMM profile.

### Completion of *L. boulardi* and *L. heterotoma* venom RhoGAP sequences

Venom apparatus, corresponding to venom glands and associated reservoirs, were dissected in Ringer’s saline. Total RNA was extracted from venom apparatus using the RNeasy Plus Micro Kit (QIAGEN) according to the manufacturers’ instructions. To obtain the full-length coding sequence of LbmGAP1.3, LbmGAP5, LbyGAP6, LhGAP1 and LhGAP2, rapid amplification of cDNA ends (RACE) was performed using the SMART RACE cDNA Amplification Kit (Clontech). For LhGAP3, 3’ RACE could not be applied due to the presence of poly(A) stretches in the sequence. LhGAP3-matching sequences were searched for in the transcriptome assembly obtained by Goecks et al., 2013) from the abdomen of *L. heterotoma* females (GenBank accession number GAJC00000000) using the command line NCBI-BLAST package (version 2.2.24). The resulting complete LhGAP3 sequence was then verified by RT-PCR using the iScript cDNA Synthesis Kit (BioRad) and the GoTaq DNA Polymerase (Promega), followed by direct sequencing of the amplified fragment. Geneious software (Biomatters) was used for sequence editing and assembly.

### Obtention of *Leptopilina* RacGAP1 coding sequences

A combination of RT-PCR and RACE using conserved and specific primers was used to clone the full-length ORF sequence of *L. boulardi* ISm and *L. heterotoma* RacGAP1. The amino acid sequence of *Nasonia vitripennis* RacGAP1 (Supplementary Table S2) was used to search for similar sequences among Hymenoptera species using BLASTP at NCBI (http://www.ncbi.nlm.nih.gov/blast/). A multiple sequence alignment of the RacGAP1 amino acid sequences found in Hymenoptera was then performed using MUSCLE (Edgar 2004). Two pairs of RacGAP1-specific degenerate primers were designed from the identified conserved regions (Supplementary Figure S1). Total RNA was extracted from Lbm and Lh individuals using the TRIzol reagent (Invitrogen), and RT-PCR experiments were performed using the RacGAP1-specific degenerate primers. After direct sequencing of the amplified fragments, Lbm- and Lh-specific primers were designed to complete the sequences obtained by RT-PCR and RACE (Supplementary Figure S1). The coding sequence of *L. clavipes* RacGAP1 was obtained by BLAST searches in the corresponding transcriptome using *L. boulardi* ISm and *L. heterotoma* RacGAP1 sequences as queries.

### Obtention of the genomic sequences of *Leptopilina* venom RhoGAPs and RacGAP1

The genomic sequences of *Leptopilina* venom RhoGAPs and RacGAP1 were obtained by BLAST and Exonerate searches (Slater & Birney 2005) using the corresponding coding sequences as queries. *L. boulardi* ISm, *L. boulardi* ISy, *L. heterotoma* and *L. clavipes* genome assemblies were obtained from NCBI under accession numbers GCA_011634795.1, GCA_019393585.1, GCF_015476425.1 and GCA_001855655.1 respectively.

### Search for candidate *V. canescens* RhoGAP calyx sequences

Coding regions were identified from the *V. canescens* calyx transcriptome published by Pichon et al., (2015) and translated into protein sequences using TransDecoder (Haas et al., 2013). The search for RhoGAP domain-containing sequences was performed using hmmsearch from the HMMER package (Eddy 2009) with the RhoGAP (PF00620) HMM profile. The identified RhoGAP calyx sequences were used as a database for Mascot to explore MS/MS data obtained from purified *V. canescens* VLPs (Pichon et al., 2015) with a false discovery rate of 1%. Only proteins identified by two or more peptides were considered.

The *V. Canescens* RacGAP1 coding sequence was obtained by BLAST searches in the *V. canescens* calyx transcriptome using the *N. vitripennis* RacGAP1 sequence as query.

The *V. canescens* calyx RhoGAP and RacGAP1 genome sequences were obtained by BLAST and Exonerate searches (Slater and Birney 2005) using the corresponding coding sequences as queries. The *V. canescens* genome assembly was obtained from NCBI under accession number GCA_019457755.1.

### Identification of domains and motifs and prediction of subcellular localization

The presence and position of signal peptide cleavage sites in the identified RhoGAP domain-containing sequences were predicted using the SignalP server (https://services.healthtech.dtu.dk/service.php?SignalP). The searches for domains and motifs were performed using InterProScan 5 (Jones et al., 2014) on the InterPro integrative protein signature database (https://www.ebi.ac.uk/interpro/). Coiled-coil regions were predicted using COILS (Lucas et al., 1991).

To identify the possible origin of the signal peptide of *Leptopilina* venom RhoGAPs, we performed a similarity search using Exonerate (Slater and Birney 2005) with the exon(s) preceding the RhoGAP domain in the genomes of *L. boulardi* ISm and ISy, *L. clavipes* and *L. heterotoma*.

Prediction of protein subcellular localization and sorting signals for *Venturia* RacGAP1 and calyx RhoGAPs was performed using the DeepLoc2 server (https://services.healthtech.dtu.dk/services/DeepLoc-2.0/).

Nuclear localization for VcRacGAP1 and calyx RhoGAPs was also predicted using the following three tools: NucPred (Brameier et al., 2007) was used to indicate whether a protein spends part of its time in the nucleus (https://nucpred.bioinfo.se/nucpred/), LocTree3 (Goldberg et al., 2014) to provide subcellular localization and Gene Ontology terms (https://rostlab.org/services/loctree3/), and PSORT II (Nakai & Horton 1999) to detect sorting signals and subcellular localization (https://psort.hgc.jp/form2.html). PSORT II results corresponding to ER membrane retention signals and cleavage sites for mitochondrial presequences were included as they may be associated with nuclear localization. Mitochondrial presequences have no sequence homology but possess physical characteristics that allow them to interact with the outer membranes of mitochondria and thus allow the targeting of proteins to the mitochondrial matrix. Regarding mitochondrial targeting, it has been shown from plants to humans that some proteins with mitochondrial presequences are dually targeted to mitochondria and the nucleus (Millar et al., 2006; Mueller et al., 2004). Indeed, anchoring signals to the ER membrane are known to allow baculoviral proteins to migrate to the inner nuclear membrane (Braunagel et al., 1996). Proteins forming VLPs in *V. canescens* could thus follow the same pathways as their homologous proteins in baculoviruses, nudiviruses, hytrosaviruses, and bracoviruses due to the conservation of this mechanism among these viruses belonging to the order Lefavirales (Braunagel et al., 1996; Hong et al., 1997; Braunagel et al., 2004; Abd-Alla et al., 2008; Bézier et al., 2009; Braunagel et al., 2009).

### Phylogeny of *Leptopilina* venom and *Venturia* calyx RhoGAPs

Searches for *N. vitripennis* RhoGAP sequences were performed using BLASTP at NCBI (http://www.ncbi.nlm.nih.gov/blast/) and hmmsearch from the HMMER package (Eddy 2009) with the RhoGAP (PF00620) HMM profile on the *N. vitripennis* v2 proteome database (Rago et al., 2016).

For the phylogenetic analysis of *Leptopilina* venom and *Venturia* calyx RhoGAPs, multiple alignments of the RhoGAP domain amino acid sequences were performed using MAFFT with the--auto option (Katoh and Standley 2013). For codon-based analysis of selection (see below), codon-based alignments of complete coding sequences were performed using RevTrans (https://services.healthtech.dtu.dk/service.php?RevTrans) with the amino acid alignments as templates. Poorly aligned regions were removed using trimAl with the - automated1 option (Capella-Gutierrez et al., 2009). Phylogenetic analyses were performed using maximum likelihood (ML) with IQ-TREE (Minh et al., 2020). ModelFinder was used to select the best model selection for phylogeny (Kalyaanamoorthy et al., 2017). The alignments obtained before and after using Trimal are shown in Supplementary Dataset 1.

### Codon-based analysis of selection

Codon-based alignments of complete coding sequences were performed with RevTrans using the amino acid alignments as templates (see above). Phylogenetic analyses were performed using maximum likelihood (ML) with PhyML (Guindon et al., 2010). Smart Model Selection was used to choose the best model selection for the phylogeny (Lefort et al., 2017).

To detect evolutionary selective pressures acting on RhoGAP sequences, the ratios of non-synonymous substitutions (dN) to synonymous substitutions (dS) were compared using different ML frameworks: the CODEML program in the PAML (Phylogenetic Analysis by Maximum Likelihood) package (Yang 1997; Yang 2007), the HyPhy software implemented at http://www.datamonkey.org/ (Delport et al., 2010), and the Selecton server (Stern et al., 2007) available at http://selecton.tau.ac.il/index.html.

Five different methods were used to detect codons under positive selection. In the first, codon substitution models implemented in CodeML were applied to the codon-based alignment using the F3×4 frequency model. Two pairs of site models were used to determine whether some codons were under positive selection: M1a (neutral) versus M2a (selection) and M7 (beta distribution where 0 < ω (dN/dS ratio) *<* 1) versus M8 (beta distribution as in M7 with ω > 1 as additional class). The models were compared using a likelihood ratio test (LRT) with 2 degrees of freedom to assess the significance of detection of selection (Yang et al., 2000). Bayes Empirical Bayes (BEB) inference was then used to identify amino acid sites with a posterior probability >95% of being under positive selection (Yang et al., 2005). We then applied four other methods of detection of selection available in the HyPhy package: Single Likelihood Ancestor Counting (SLAC), Fixed Effect Likelihood (FEL), Mixed Effects Model of Evolution (MEME) and Fast Unbiased Bayesian AppRoximation (FUBAR) methods (Kosakovsky Pond & Frost 2005; Murrell et al., 2012; Murrell et al., 2013). To eliminate false-positive detection, only codons identified by CodeML M2a and M8 and at least one of the other methods were considered under positive selection. Radical or conservative replacements were then determined based on whether they involved a change in the physicochemical properties of a given amino acid, such as charge or polarity (Zhang 2000).

Four different methods were used to detect codons under negative selection: the Selecton server using the pair of site models M8a (ß & ω=1) versus M8 (ß & ω), and the FEL, SLAC and FUBAR methods mentioned above. Only codons identified by Selecton and at least one other method were considered under purifying selection.

To identify specific lineages with a proportion of sites evolving under positive or purifying selection, we performed branch-site REL analyses using the HyPhy package (Kosakovsky Pond et al., 2011). Unlike the branch and branch-site lineage-specific models available in CodeML, branch-site REL does not require *a priori* identification of foreground and background branches.

### Structural analysis

Molecular modeling of VcVLP2 RhoGAP was performed using the Phyre server with default parameters (http://www.sbg.bio.ic.ac.uk/phyre2/) (Kelley et al., 2015). The model with the highest confidence (100%) and coverage (46%) was obtained using the crystal structure with PDB code 2OVJ as a template. Model quality was evaluated using the QMEAN server (http://swissmodel.expasy.org/qmean/) (Benkert et al., 2011). Briefly, the QMEAN score is a global reliability score with values ranging between 0 (lower accuracy) and 1 (higher accuracy). The associated Z-score relates this QMEAN score to the scores of a non-redundant set of high-resolution X-ray structures of similar size, with ideal values close to 0. Visualization of LbmGAP (Colinet et al., 2007) and VcVLP2 structures and mapping of sites under selection were performed using PyMol (http://sourceforge.net/projects/pymol/). Secondary structure assignment was performed with the DSSP program (Kabsch & Sander 1983). Accessible surface area (ASA) or solvent accessibility of amino acids was predicted using the *ASAView* algorithm (Ahmad et al., 2004).

### *In vitro* mutagenesis

The S76K, V124K, L137D, T143L, S150L, I185K, E200G, and V203R mutations were introduced into the LbmGAP cDNA using the QuickChange XL Site-Directed Mutagenesis Kit (Stratagene). The results of *in vitro* mutagenesis were verified by sequencing.

### Yeast two-hybrid analysis

Interactions between LbmGAP and LbmGAP mutants and mutated forms of *Drosophila* RhoA, Rac1, Rac2 and Cdc42 GTPases were individually examined by mating as previously described (Colinet et al., 2007). The plasmids expressing the GTPase proteins were tested against the pGADT7-T control vector, which encodes a fusion between the GAL4 activation domain and the SV40 large T-antigen. Reciprocally, plasmids producing LbmGAP and LbmGAP mutants were tested against the pLex-Lamin control vector. Interactions between LbmGAP and the Rac1 and Rac2 GTPases were used as positive controls (Colinet et al., 2007). Interactions were initially tested by spotting five-fold serial dilutions of cells on minimal medium lacking histidine and supplemented with 3-amino-triazole at 0.5 mM to reduce the number of false positives. ß-galactosidase activity was then detected on plates (Fromont-Racine et al., 1997).

### Purification of *Leptopilina* venosomes and mass spectrometry analysis

Twenty-five Lbm, Lby and Lh female venom apparatus were dissected and the reservoirs separated from the gland. The twenty-five reservoirs were then pooled and opened to release the venom content in 25 µl of Ringer’s solution supplemented with a protease inhibitor cocktail (Sigma). The venom suspension was centrifuged at 500xg for 5 min to remove the residual tissues, then centrifuged at 15,000xg for 10 min to separate the vesicular fraction from the soluble venom proteins (supernatant fraction) (Wan et al., 2019). The vesicular fraction was then washed twice by resuspension in 25 µl of Ringer’s solution followed by centrifugation at 15,000xg for 10 min. The two samples were mixed with 4x Laemmli buffer containing ß-mercaptoethanol (v/v) and boiled for 5 min. Proteins were then separated on a 6–16% linear gradient SDS-PAGE and the gel was silver stained as previously described (Colinet et al., 2013). Identification of proteins by nano-LC-tandem mass spectrometry (MS/MS) was performed on bands excised from the gels as previously described (Colinet et al., 2013). MS/MS data analysis was performed with Mascot software (http://www.matrixscience.com), licensed in house using the full-length coding sequences of the *Leptopilina* venom RhoGAP sequences. Data validation criteria were (i) one peptide with individual ion score greater than 50 or (ii) at least two peptides of individual ion score greater than 20. The mascot score is calculated as −10Log (p). The number of peptide matches identified by Mascot software on the MS/MS data was used to determine (i) the number of protein bands in which each venom RhoGAP was found and (ii) the protein band in which each venom RhoGAP was the most abundant. The calculated FDR (based on an automated decoy database search) was less than 1%. The mass spectrometry proteomics data have been deposited to the ProteomeXchange Consortium via the PRIDE (Perez-Riverol et al., 2022) partner repository with the dataset identifier PXD041695.

## Results

### *Leptopilina* venom RhoGAPs probably emerged in the ancestor of the genus

In the Figitidae, venom RhoGAPs have only been described for *L. boulardi* and *L. heterotoma* (Labrosse et al., 2005; Colinet et al., 2013). A search for candidate homologs was performed in the transcriptomes of *L. victoriae* and *L. clavipes*, two other *Leptopilina* species, *Gasnaspis* sp. G1, another member of the family Figitidae, and *Andricus quercuscalicis* and *Synergus umbraculus,* two gall wasp species representatives of the superfamily Cynipoidea outside of Figitidae (separated by approximately 50 Mya (Peters et al., 2017)). The putative venom RhoGAPs were identified based on the presence of a signal peptide at the N-terminus of the protein, followed by a RhoGAP domain (Table 1). Two transcript sequences encoding RhoGAP domain-containing proteins predicted to be secreted were found in *L. victoriae* (LvGAP1 and LvGAP2) and one in *L. clavipes* (LcGAP1), whereas none were found for *Gasnaspis* sp. G1, *A. quercuscalicis* and *S. umbraculus* (Table 1 and Figure 1A). Our results therefore suggest that venom RhoGAPs are specific to the genus *Leptopilina* although further taxon sampling would be required to fully support this hypothesis.

**Figure 1.**
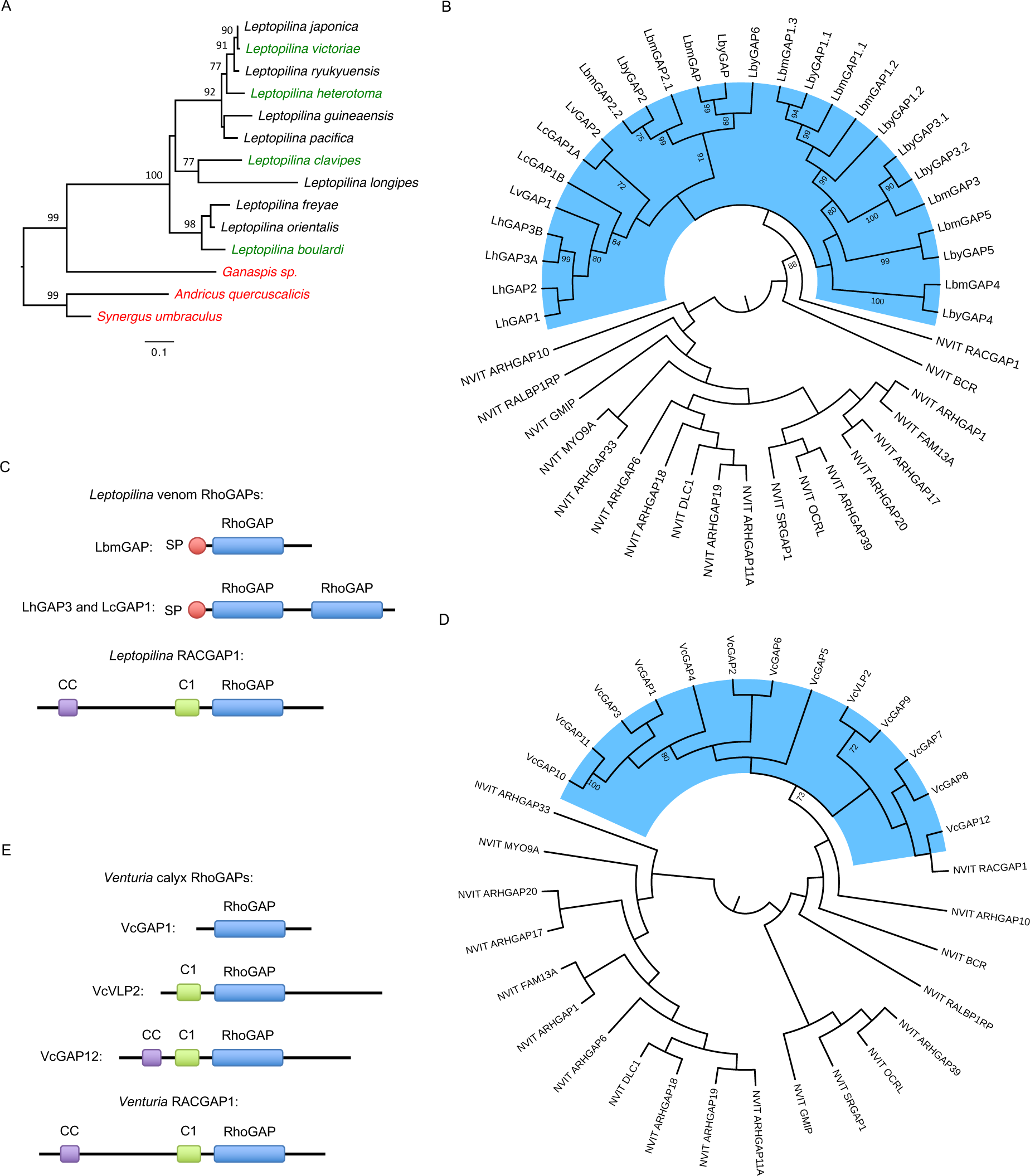
Origin of *Leptopilina* venom RhoGAPs and *Venturia* calyx RhoGAPs. (A) Maximum-likelihood phylogenetic tree of selected members of the Cynipoidea superfamily. The phylogenetic tree was obtained with IQ-TREE using internal transcribed spacer 2 (ITS2) sequence. Species in red are those for which venom RhoGAPs were not found. No data were available for species in black. The tree was rooted on the branch that leads to the two gall wasp species, *Andricus quercuscalicis* and *Synergus umbraculus.* The scale bar indicates the number of substitutions per site. (B) Maximum-likelihood phylogenetic tree of *Leptopilina* venom RhoGAPs along with *N. vitripennis* classical RhoGAPs. (C) Comparison of protein domain organization between LbmGAP, representative of most *Leptopilina* venom RhoGAPs, LhGAP3 and LcGAP1, the two venom RhoGAPs that contain two RhoGAP domains, and *Leptopilina* RacGAP1. (D) Maximum-likelihood phylogenetic tree of *Venturia* calyx RhoGAPs along with *N. vitripennis* classical RhoGAPs. (E) Comparison of protein domain organization between VcGAP1, VcVLP2 and VcGAP12, representative of the other *Venturia* calyx RhoGAPs, and *Venturia* RacGAP1. In A, B, and D, numbers at corresponding nodes are bootstrap support values in percent (500 bootstrap replicates). Only bootstrap values greater than 70% are shown. In B and D, the phylogenetic tree was obtained with IQ-TREE using the RhoGAP domain amino acid sequence and displayed as a cladogram. *Leptopilina* venom RhoGAPs and *Venturia* calyx RhoGAPs are highlighted in blue. The tree was rooted at midpoint. In C and E, C1: protein kinase C-like zinc finger motif, CC: coiled-coil region, RhoGAP: RhoGAP domain found in RacGAP1-related proteins, SP: signal peptide.

**Table 1.**
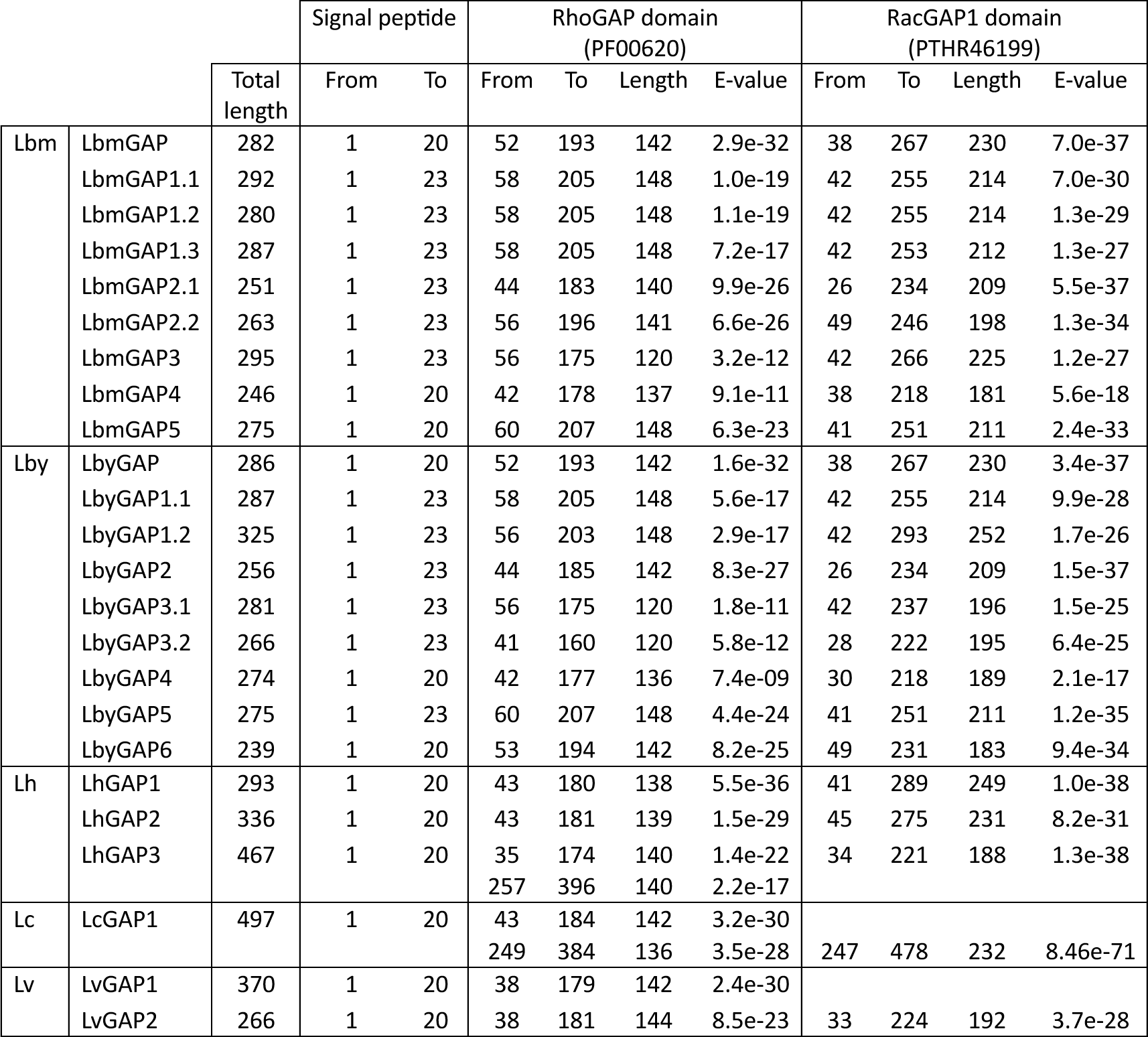
Motif and domain organization of *Leptopilina* venom RhoGAPs. The signal peptide was predicted using SignalP at CBS. The RhoGAP domain (PF00620) from the Pfam database and the RacGAP1 domain (PTHR46199) from the Panther database were identified using InterProScan on the InterPro integrative protein signature database. Compared to Pfam, the Panther database provides information about the specific RhoGAP subfamily to which the domain belongs. Two successive RhoGAP domains were found in LhGAP3 and LcGAP1. Lbm: *L. boulardi* ISm; Lby: *L. boulardi* ISy; Lh: *L. heterotoma*; Lc: *L. clavipes*; Lv: *L. victoriae.* The dot numbering (e.g., LbmGAP1.1, LbmGAP1.2, and LbmGAP1.3) was used for *Leptopilina* venom RhoGAPs that are encoded by different genes but share high amino acid sequence identity in the RhoGAP domain (greater than 70% identity).

### *Leptopilina* venom RhoGAPs likely evolved from a single imperfect duplication of RacGAP1

A phylogenetic analysis was performed to learn more about the evolutionary history of *Leptopilina* venom RhoGAPs. The phylogeny was constructed based on the amino acid sequence of the RhoGAP domain of these venom proteins and that of the 19 RhoGAPs we identified from the predicted proteome of the jewel wasp *Nasonia vitripennis* (Supplementary Table S2)*. N. vitripennis* (superfamily Chalcidoidea) was chosen because it was the first parasitoid wasp to have its genome sequenced, and its genome has recently been re-annotated (Rago et al., 2016). Furthermore, the only RhoGAPs found in *N. vitripennis* are classical intracellular RhoGAPs, which are absent in maternal fluids (de Graaf et al., 2010).

Prior to phylogenetic analysis, the complete coding sequence was obtained for two Lbm (LbmGAP1.3 and LbmGAP5), one Lby (LbyGAP6) and all three Lh venom RhoGAPs as described in the Methods section. Two of the *Leptopilina* venom RhoGAPs, namely LhGAP3 and LcGAP1, were predicted to contain two RhoGAP domains instead of one. Therefore, these two domains were separated for the analysis. The resulting phylogeny identified NvRacGAP1 as the closest RhoGAP from *N. vitripennis* to all *Leptopilina* venom RhoGAPs with confident support values (Figure 1B). Accordingly, the RhoGAP domain found in all *Leptopilina* venom RhoGAPs was predicted to belong to the RacGAP1 subfamily typically found in RacGAP1-related proteins (Table 1). These results suggest that the *Leptopilina* venom RhoGAPs derive from duplication of the *N. vitripennis* RacGAP1 ortholog in *Leptopilina*.

The RacGAP1 coding sequence of *L. boulardi* ISm*, L. heterotoma and L. clavipes* was obtained either by cloning and sequencing or by searching genomic data, and its domain organization was compared with that of venom RhoGAPs (Figure 1C, Table 1 and Supplementary Figure S2). *Leptopilina* RacGAP1 has a typical domain organization consisting of a coiled-coil region and a C1 motif (protein kinase C-like zinc finger motif, a ∼ 50 amino-acid cysteine-rich domain involved in phorbol ester/diacylglycerol and Zn^2+^ binding) followed by a RhoGAP domain (Tcherkezian & Lamarche-Vane 2007). A large part of the RacGAP1 sequence spanning the coiled-coil region and the C1 motif is replaced by the signal peptide in all *Leptopilina* venom RhoGAPs (Figure 1C, Table 1 and Supplementary Figure S2). This suggests that a unique incomplete duplication of the RacGAP1 gene occurred in the ancestor of the *Leptopilina* genus, resulting in a truncated duplicate encoding a RhoGAP with an altered N-terminus.

To further investigate this duplication event and the possible origin of the signal peptide, the genomic sequences of *L. boulardi* ISm, *L. heterotoma* and *L. clavipes* venom RhoGAPs and RacGAP1 were obtained, either by cloning and sequencing or by searching in genomic data, and compared (Supplementary Table S3). The genomic sequence of *Leptopilina* RacGAP1 consists of 10 exons with a “CC region”, “C1 motif” and “RhoGAP domain” encoded by exons 2 and 3, exons 5 and 6 and exons 7 and 8, respectively. The RhoGAP domain is also encoded by two exons for venom RhoGAPs. These two exons are preceded by one to three exons, with the signal peptide encoded by the first exon for most of them. Sequence comparisons revealed similarities between venom RhoGAPs and *Leptopilina* RacGAP1 only for the sequence spanning the two exons encoding the RhoGAP domain and part of the following one. No significant similarities were found for the preceding exons at either the nucleotide or amino acid level (Supplementary Figure S3). This supports the hypothesis of a partial duplication of the RacGAP1 gene in the ancestor of the *Leptopilina* genus, resulting in the loss of the sequence spanning exons 1 to 6, followed by further duplication of the ancestrally duplicated gene during the diversification of the genus *Leptopilina*. No significant sequence similarity was found between the exon(s) preceding the RhoGAP domain coding sequence of *Leptopilina* venom RhoGAPs and other sequences in the genomes of *L. boulardi* ISm and ISy, *L. clavipes*, and *L. heterotoma* that could provide an indication of the origin of the signal peptide (Supplementary Dataset 2).

Some of the duplication events that followed the initial duplication of the RacGAP1 gene in the ancestor of the *Leptopilina* genus appear to be specific to *L. boulardi* and would explain the large number of venom RhoGAPs found in this species (Figure 1B and Supplementary Figure S2). The two consecutive RhoGAP domains found in LhGAP3 were grouped together with confident support values in the phylogenies (Figure 1B and Supplementary Figure S2). This suggests that a partial tandem duplication spanning exons 2 and 3 has occurred for this *L. heterotoma* venom RhoGAP. In contrast, the two RhoGAP domains found in LcGAP1 did not group together (Figure 1B and Supplementary Figure S2). However, the genomic sequence of LcGAP1 was not complete and we could not find the region corresponding to the second RhoGAP domain in the *L. clavipes* genome (Supplementary Table S3). Therefore, it is unclear whether the LcGAP1 coding sequence found in the transcriptome assembly is true or artifactual.

### *L. boulardi* and *L. heterotoma* venom RhoGAPs are associated with venosomes

LbmGAP and LbmGAP2 have been shown to be associated with vesicles named venosomes produced in *L. boulardi* venom that transport them to *Drosophila* lamellocytes (Wan et al., 2019). Our next goal was to investigate whether *Leptopilina* venom RhoGAPs other than LbmGAP and LbmGAP2 could be associated with venosomes. Proteomic analysis was performed on the supernatant and vesicular fractions separated from the venom by centrifugation. Comparison of the electrophoretic profiles obtained on a 6-16% SDS-PAGE for *L. boulardi* ISm (Lbm), *L. boulardi* ISy (Lby) and *L. heterotoma* (Lh) revealed an important variation between both fractions for all three wasps (Figure 2). All the major bands on the electrophoretic patterns of Lbm, Lby and Lh supernatant and vesicular fractions, as well as several minor bands (35 bands in total for Lbm, 34 for Lby and 37 for Lh), were excised and tryptic peptides were analyzed by mass spectrometry. The coding sequences of *Leptopilina* venom RhoGAPs were used to perform Mascot searches on the mass spectrometry data obtained from both venom fractions. All *L. boulardi* and *L. heterotoma* venom RhoGAPs were detected in the vesicular fraction, where most of them were found to be enriched (Figure 2). We could then hypothesize that they are associated with and transported by venosomes to target *Drosophila* cells.

**Figure 2.**
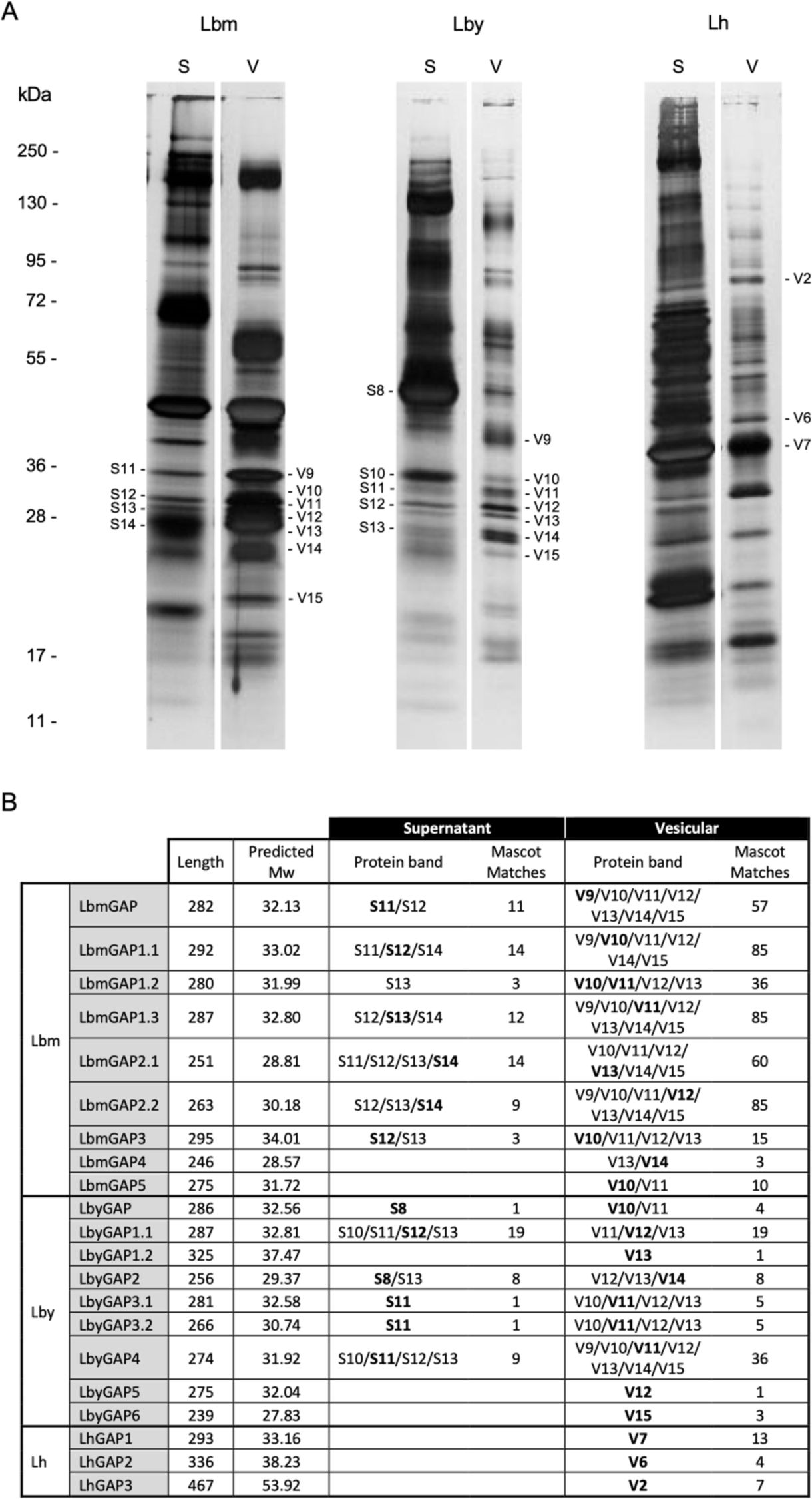
Proteomic analysis of *Leptopilina* venom supernatant and vesicular fractions. (A) The supernatant (S) and vesicular (V) fractions obtained from 25 Lbm, Lby and Lh venom reservoirs were separated on a 6-16% SDS-PAGE under reducing conditions and visualized by silver staining. Protein bands in which *Leptopilina* venom RhoGAPs were identified by mass spectrometry are numbered on the gel. Molecular weight standard positions are indicated on the left (kDa). (B) For each *Leptopilina* venom RhoGAP, length (number of amino acids), predicted Mw (kDa), bands in which specific peptides were found by mass spectrometry and number of total Mascot matches using the coding sequences of *Leptopilina* venom RhoGAPs as database and according to the S and V venom fractions are given. Numbers in bold correspond to the bands in which each RhoGAP was identified as the most abundant according to the number of Mascot matches.

### *V. canescens* calyx RhoGAPs probably evolved from two or more imperfect RacGAP1 duplication events

Analysis of the *V. canescens* calyx transcriptome allowed the identification of a total of 13 RhoGAP domain-containing protein coding sequences, including VcVLP2 (Table 2 and Supplementary Table S4). Matches with peptides obtained from a proteomic analysis of *V. canescens* VLPs (Pichon et al., 2015) were detected for all 13 calyx RhoGAPs, indicating that they are associated with VLPs (Table 2).

**Table 2.** Motif and domain organization of *Venturia* calyx RhoGAP proteins associated with VLPs. The coiled-coil (CC) region was predicted using COILS (Lucas et al., 1991). The protein kinase C-like zinc finger (C1) motif (PS50081) from the Prosite database, the RhoGAP domain (PF00620) from the Pfam database and the RacGAP1 domain (PTHR46199) from the Panther database were identified using InterProScan on the InterPro integrative protein signature database. Compared to Pfam, the Panther database provides information about the specific RhoGAP subfamily to which the domain belongs. Mascot: number of matches with peptides from purified *V. canescens* VLPs. ^a^Coding sequence not complete at 5’ end. ^b^Coding sequence not complete at 3’ end.

|  | Mascot | Total length | CC |  | C1 (PS50081) |  | RhoGAP domain (PF00620) |  |  |  | RacGAP1 domain (PTHR46199) |  |  |  |
| --- | --- | --- | --- | --- | --- | --- | --- | --- | --- | --- | --- | --- | --- | --- |
|  |  |  | From | To | From | To | From | To | Length | E-value | From | To | Length | E-value |
| VcVLP2 | 26 | 485 |  |  | 56 | 106 | 134 | 277 | 144 | 7.4e-36 | 28 | 366 | 339 | 3.1e-98 |
| VcGAP1 | 9 | 207 |  |  |  |  | 30 | 168 | 139 | 5.7e-25 | 12 | 199 | 188 | 5.1e-49 |
| VcGAP2 | 19 | 280 |  |  |  |  | 23 | 161 | 139 | 6.9e-29 | 8 | 195 | 188 | 1.5e-50 |
| VcGAP3 | 4 | 270 |  |  | 22 | 69 | 92 | 231 | 140 | 2.6e-25 | 14 | 265 | 252 | 2.9e-63 |
| VcGAP4 | 21 | 382 |  |  | 37 | 86 | 116 | 240 | 125 | 2.2e-16 | 20 | 298 | 279 | 2.7e-68 |
| VcGAP5 | 19 | 307 |  |  | 15 | 63 | 91 | 236 | 146 | 4.0e-24 | 7 | 281 | 275 | 2.6e-60 |
| VcGAP6 | 16 | 284 |  |  | 17 | 67 | 88 | 215 | 128 | 3.3e-19 | 14 | 270 | 257 | 8.0e-55 |
| VcGAP7 | 14 | 225<sup>a</sup> |  |  |  |  | 15 | 156 | 142 | 5.3e-29 | 4 | 212 | 209 | 9.2e-57 |
| VcGAP8 | 5 | 369 |  |  | 64 | 113 | 141 | 281 | 141 | 4.6e-28 | 20 | 334 | 315 | 3.6e-84 |
| VcGAP9 | 9 | 319<sup>b</sup> |  |  | 54 | 104 | 123 | 266 | 144 | 1.1e-28 | 38 | 306 | 269 | 5.5e-73 |
| VcGAP10 | 5 | 233 |  |  |  |  | 30 | 166 | 137 | 1.6e-22 | 7 | 219 | 213 | 7.1e-45 |
| VcGAP11 | 5 | 199 |  |  |  |  | 27 | 166 | 140 | 4.3e-19 | 8 | 198 | 191 | 5.7e-42 |
| VcGAP12 | 8 | 402 | 21 | 48 | 79 | 131 | 159 | 296 | 138 | 3.3e-35 | 54 | 372 | 319 | 9.7e-91 |

Similar to the *Leptopilina* venom RhoGAPs, the *V. canescens* calyx RhoGAPs contain a RacGAP1 domain (Table 2) and are closely related to NvRacGAP1 (Figure 1D). A phylogenetic analysis was performed on *Leptopilina* venom RhoGAPs and *V. canescens* calyx RhoGAPs, together with *Leptopilina* and *V. canescens* RacGAP1 and *N. vitripennis* classical RhoGAPs, using either protein-or codon-based alignment of the RhoGAP domain (Supplementary Figure S4). The analysis confirmed a relationship between either *Leptopilina* venom RhoGAPs or *V. canescens* calyx RhoGAPs with RacGAP1. In the codon-based phylogeny, *V. canescens* calyx RhoGAPs formed a robust monophyletic group with VcRacGAP1, suggesting an independent duplication event (Supplementary Figure S4). However, in the protein-based phylogeny, the *V. canescens* calyx RhoGAPs did not form a robust monophyletic group with VcRacGAP1. Similarly, the *Leptopilina* venom RhoGAPs did not group with *Leptopilina* RacGAP1 in either phylogeny. This is likely due to the high divergence of *Leptopilina* venom RhoGAPs and *V. canescens* calyx RhoGAPs compared to RacGAP1 (Supplementary Figure S4).

In contrast to the *Leptopilina* venom RhoGAPs, none of the *V. canescens* calyx proteins were predicted to be secreted. Most were predicted to be localized in the nucleus and/or to contain a nuclear localization signal (Supplementary Tables S5 and S6). The absence of a predicted nuclear localization signal for VcGAP7 may be explained by the incomplete coding sequence at the 5’ end (Table 2). There is greater variation in domain organization for *V. canescens* calyx RhoGAPs compared to *Leptopilina* venom RhoGAPs (Figure 1E and Supplementary Figure S5). Eight *V. canescens* calyx RhoGAPs have retained the C1 motif. In addition, VcGAP12 still contains a CC region in the N-terminal part of the sequence although the total sequence length is shorter than RacGAP1 (Figure 1E and Supplementary Figure S5).

The genomic coding sequence of VcRacGAP1 was obtained by genomic data mining. It comprised 9 exons with the CC region, C1 motif and RhoGAP domain encoded by exons 2 and 3, exons 5 and 6 and exon 7, respectively (Supplementary Table S7). In contrast to *Leptopilina*, the RhoGAP domain is encoded by a single exon in *V. canescens* RacGAP1 as well as in all calyx RhoGAPs. Interestingly, the VcGAP12 gene is located close to the VcRacGAP1 gene on chromosome 3, but is three exons shorter, suggesting a recent incomplete duplication (Supplementary Table S7). However, no phylogenetic relationship could be inferred between VcRacGAP1 and VcGAP12, as the corresponding proteins were either grouped in the phylogeny but the grouping was not supported by bootstrap analysis (Figure 1D), or were not grouped in the phylogeny (Supplementary Figures S4 and S5). In contrast to VcRacGAP1 and VcGAP12, VcVLP2 and the other VcGAP genes are more widely distributed in the *V. canescens* genome, being organized into several clusters consisting of two or more genes in tandem arrays (Figure 3 and Supplementary Table S7). For several of these clusters, the *V. canescens* calyx RhoGAP proteins were found in bootstrap-supported groups in the phylogeny, suggesting a close relationship (Figure 1D, Supplementary Figure S5 and Figure 3). This is the case, for example, for VcVLP2 and VcGAP9, VcGAP1 and VcGAP3, or VcGAP10 and VcGAP11, which most likely originated from tandem duplication events. Therefore, it is possible that two or more incomplete duplication events of the VcRacGAP1 gene, followed by further tandem and dispersed duplication of the ancestrally duplicated genes, led to the current number of calyx RhoGAPs in *V. canescens*.

**Figure 3.**
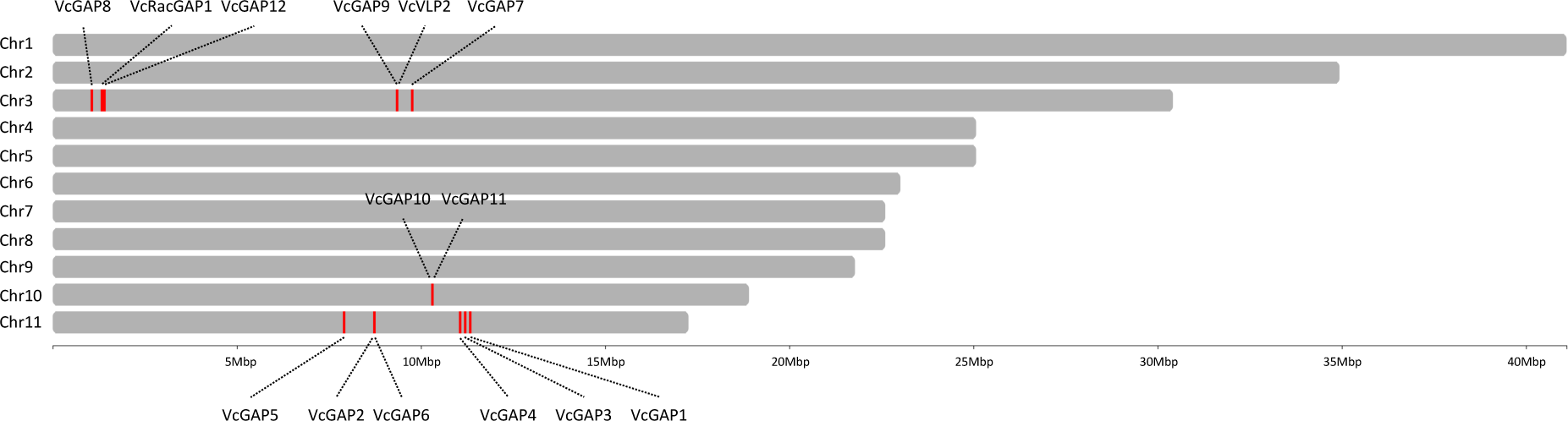
*V. canescens* chromosome map with the position of gene loci corresponding to RacGAP1 and calyx RhoGAPs visualized using chromoMap R package (Anand & Rodriguez Lopez 2022).

### Evidence of positive selection in *L. boulardi* venom and *V. canescens* calyx RhoGAP sequences

In a previous work, we identified four amino acid residues involved in the interaction of LbmGAP with Rho GTPases (Colinet et al., 2007), including the key arginine residue (R74 in LbmGAP) required for the GAP catalytic activity (Figure 4A). The other 8 venom RhoGAP sequences found in *L. boulardi* Ism and ISy were all mutated at this arginine residue (Figure 4B). Most also contained substitutions in one or more of the other three amino acids involved in Rho GTPase interaction (Figure 4B). In *V. canescens*, no substitutions were found for VcVLP2 at the key arginine residue or at any of the three amino acids involved in Rho GTPase interaction (Figure 4C). In contrast, all other *V. canescens* calyx RhoGAP sequences, including VcGAP9, for which a close relationship with VcVLP2 could be inferred from phylogenetic analysis and chromosomal location, exhibit substitutions in one or more of the sites essential for GAP activity and/or involved in the interaction with Rho GTPases (Figure 4C). These observations suggest that only LbmGAP (and its ortholog LbyGAP) in *L. boulardi* and VcVLP2 in *V. canescens* are functional as RhoGAPs.

**Figure 4.**
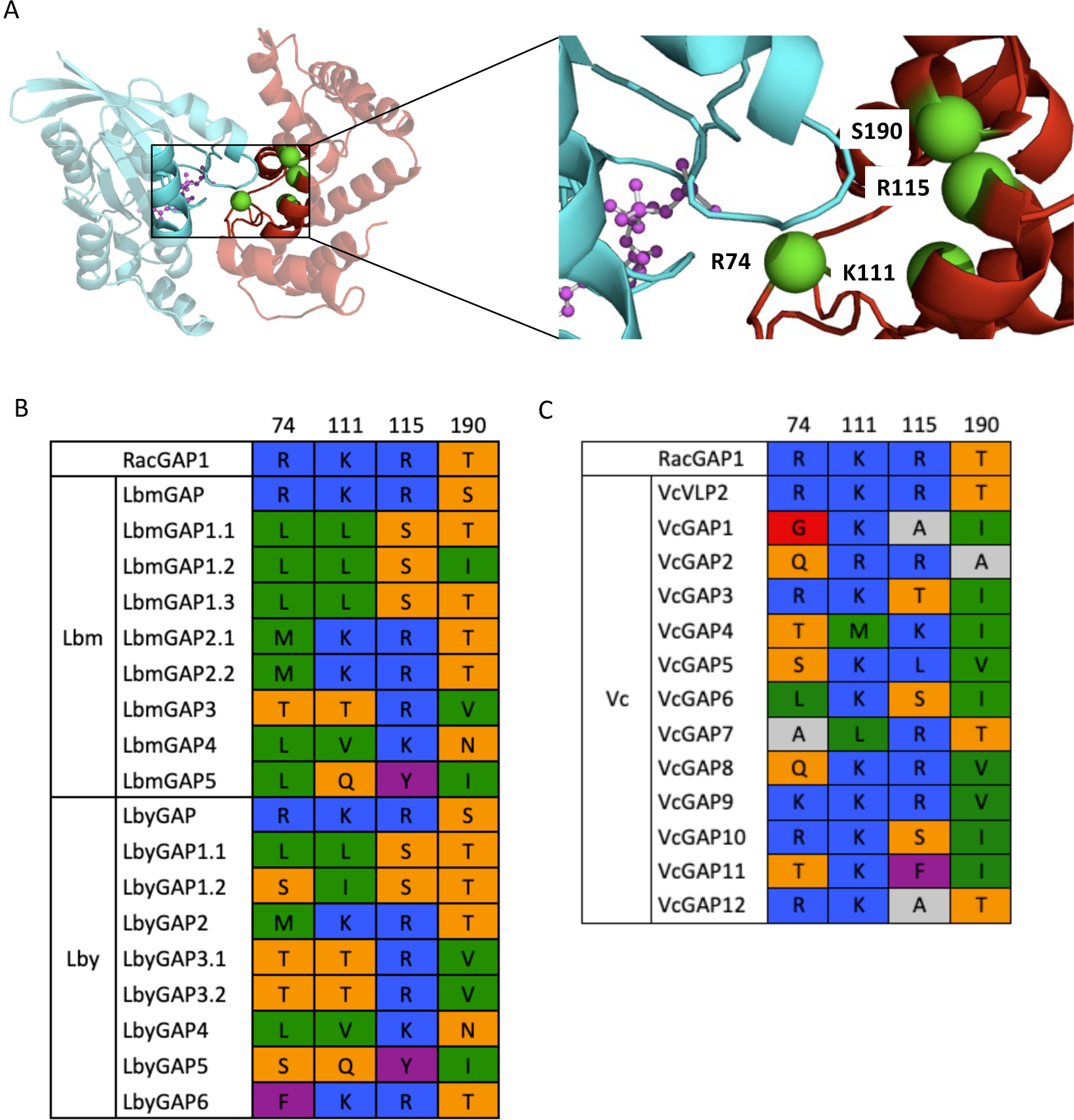
Substitutions in the essential sites for GAP activity and/or involved in interaction with Rho GTPases. (A) Tertiary structure of LbmGAP (red) in complex with Rac1 (blue) and the transition-state analogue GDP.AlF3 (modeled by homology for sequence spanning amino acid residues 51 to 216 in Colinet et al., 2007). The four sites essential for GAP activity and/or involved in interaction with RhoGTPases are colored green. (B) Amino acids found at the four sites essential for GAP activity and/or involved in the interaction with RhoGTPases for *Leptopilina* RacGAP1 and *L. boulardi* venom RhoGAP sequences. The numbering corresponds to the positions in the LbmGAP sequence. (C) Amino acids at the four sites essential for GAP activity and/or involved in interaction with RhoGTPases for *V. canescens* RacGAP1 and calyx RhoGAP sequences. The numbering corresponds to the positions in the LbmGAP sequence. In B and C, amino acids are colored according to their properties following the RasMol amino acid color scheme.

Pairwise sequence identity was lower at the amino acid level than at the nucleotide level for both *Leptopilina* venom RhoGAPs and *V. canescens* calyx RhoGAPs (Supplementary Figure S6). This indicates a higher divergence at non-synonymous sites at the codon level compared to synonymous sites. A search for positive selection at the codon level was performed to detect possible functional divergence of *L. boulardi* venom and *V. canescens* calyx RhoGAP sequences. PAML codon-based models with (M2a and M8) and without (M1a and M7) selection were compared using likelihood ratio tests (LRTs). Both M1a/M2a and M7/M8 comparisons resulted in the rejection of the null hypothesis suggesting that a fraction of codons in RhoGAP sequences are under positive selection (Table 3).

**Table 3.**
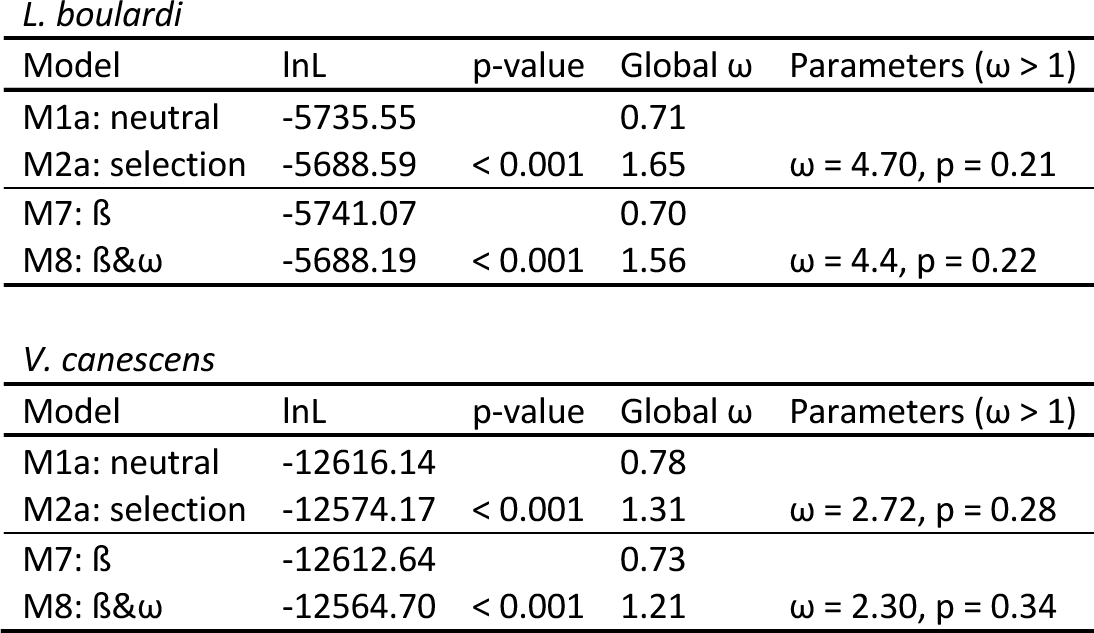
Positive selection analysis among sites using CodeML for *L. boulardi* venom and *V. canescens* calyx RhoGAP sequences. lnL is the log likelihood of the model. p-value is the result of likelihood ratio tests (LRTs). Global ω is the estimate of the dN/dS ratio under the model (given as a weighted average). Parameters (ω > 1) are parameters estimates for a dN/dS ratio greater than 1 (p is the proportion of sites with ω > 1).

Consistently, a total of 7 and 11 branches of the phylogenetic tree constructed with the *L. boulardi* venom and *V. canescens* calyx RhoGAP sequences, respectively, were detected by the REL branch-site test as corresponding to lineages on which a subset of codons has evolved under positive selection (Figure 5A and 6A). For *L. boulardi*, 6 of the 7 lineages under positive selection were internal branches, indicating that selection occurred primarily before the separation of the *L. boulardi* Ism and ISy strains (Figure 5A).

**Figure 5.**
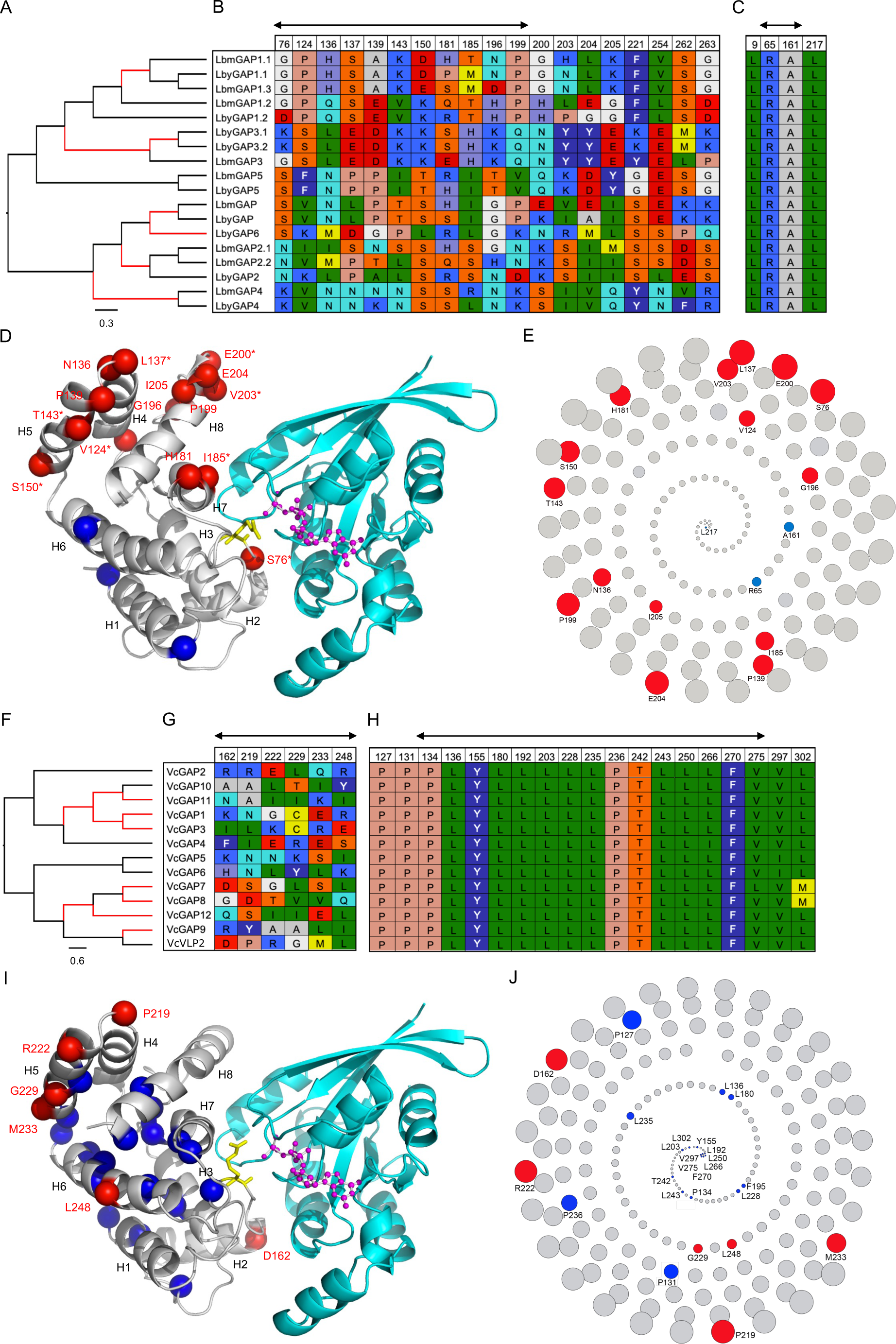
Identification of branches and codons under selection in *L. boulardi* venom RhoGAPs and *V. canescens* calyx RhoGAPs. (A) Cladogram of *L. boulardi* venom RhoGAPs. (B) Codons under positive selection numbered according to LbmGAP amino acid sequence. (C) Codons under negative selection codons numbered according to LbmGAP amino acid sequence. (D) Sites under positive and negative selection displayed on the tertiary structure of LbmGAP (grey) in complex with Rac1 (blue) and the transition-state analogue GDP.AlF3 (modeled by homology for sequence spanning amino acid residues 51 to 216 in Colinet et al., 2007). A star indicates amino acid residues tested in mutagenesis and two-hybrid interaction assays. (E) Spiral view of LbmGAP amino acids in the order of their solvent accessibility. (F) Cladogram of *V. canescens* calyx RhoGAPs. (G) Codons under positive selection numbered according to VcVLP2 amino acid sequence. (H) Codons under negative selection numbered according to VcvLP2 amino acid sequence. (I) Sites under positive and negative selection displayed on the tertiary structure of VcVLP2 (grey) in complex with Rac1 (blue) and the transition-state analogue GDP.AlF3 (modeled by homology for sequence spanning amino acid residues 117 to 316). (J) Spiral view of VcVLP2 amino acids in the order of their solvent accessibility. In A and F, Branches identified under positive selection by the HyPhy BSR method are colored in red. The cladogram was rooted at midpoint. In B, C, G, and H, amino acids are colored according to their properties following the RasMol amino acid color scheme. The two-sided arrows indicate sites located in the RhoGAP domain. In D and I, the helices are shown as ribbons and numbered according to their location in the amino acid sequence of LbmGAP (D) or VcVLP2 (I). Sites under positive selection are displayed as red-colored spheres and numbered according to the LbmGAP (D) or VcVLP2 (I) sequence. Sites under negative selection are displayed as spheres and colored in blue. Protruding LbmGAP Arg74 (D) and VcVLP2 Arg156 (I) are shown as sticks and colored in yellow. GDP.AlF3 is shown as ball-and-sticks models and colored in magenta. In E and J, Most accessible amino acids come on the outermost ring of the spiral whereas the buried amino acids are occurring in the innermost ring. Radius of the circles corresponds to the relative solvent accessibility. Red and blue colors are used for amino acids under positive and negative selection respectively. All other amino acids are in grey.

The combined use of five different methods identified a total of 19 codons as candidates under positive selection for the *L. boulardi* venom RhoGAP sequences, most of which were found in the region corresponding to the RhoGAP domain (Figure 5B). In contrast, only 4 amino acids were detected as evolving under negative selection in the *L. boulardi* venom RhoGAPs, respectively (Figure 5C). The leucine at position 9 in LbmGAP is located in the signal peptide, demonstrating the importance of this region. The other three amino acids in *L. boulardi* and most of those in *V. canescens* under negative selection are buried in the protein and are probably important for the structural stability (Figures 5D and 5E and Supplementary Figure S7A).

Interestingly, at least one radical change in charge and/or polarity of the corresponding amino acids was found for all selected codons (Figure 5B). For example, the polar uncharged serine residue at position 76 in LbmGAP was replaced by a negatively charged lysine residue in LbmGAP4 (Figure 5B). Since radical changes are more likely to modify protein function than conservative changes, this suggests that the identified non-synonymous substitutions may be adaptive. Interestingly, the corresponding amino acids are mostly exposed on the surface of the protein and therefore likely to interact with partners (Figures 5D and 5E and Supplementary Figure S7A).

We therefore generated site-specific mutants of LbmGAP for 8 of the 19 amino acids under positive selection and compared their binding capabilities to Rho GTPases. Two-hybrid analysis revealed a lack of interaction for five of the mutants, indicating that the corresponding amino acids are essential for interaction with Rac GTPases (Table 4). On the other hand, the remaining three mutants were still able to interact strongly with Rac1 and Rac2, suggesting that the corresponding amino acids are not involved in the interaction with Rac GTPases (Table 4).

**Table 4.** Summary of the results of interaction assays for the LbmGAP mutants. Results are based on growth on selective medium lacking histidine and qualitative ß-galactosidase overlay assays.-: no interaction; (+): very weak interaction; +++: strong interaction. T: SV40 large T-antigen negative control.

|  | LbmGAP | LbmGAP S76K | LbmGAP V124K | LbmGAP L137D | LbmGAP T143L | LbmGAP S150L | LbmGAP I185K | LbmGAP E200G | LbmGAP V203R | T |
| --- | --- | --- | --- | --- | --- | --- | --- | --- | --- | --- |
| Rac1 | +++ | - | - | +++ | +++ | +++ | - | - | - | - |
| Rac2 | +++ | - | - | +++ | +++ | +++ | - | - | - | - |
| CDC42 | (+) | (+) | (+) | (+) | (+) | (+) | (+) | (+) | (+) | (+) |
| RhoA | - | - | - | - | - | - | - | - | - | - |
| Lamin | - | - | - | - | - | - | - | - | - | - |

For the *V. canescens* calyx RhoGAP sequences, six amino acids were identified as candidates under positive selection and 19 under negative selection (Figures 5G and 5H). Four of the amino acids under positive selection are predicted to be exposed on the surface of the protein, indicating functional diversification in relation to interaction with partners as in *L. boulardi* (Figures 5I and 5J and Supplementary Figure S7B). Most of the amino acids under negative selection are buried within the protein and are likely to be important for structural stability (Figures 5I and 5J and Supplementary Figure S7B).

## Discussion

### Independent convergent recruitment of vesicle-associated RhoGAP proteins in distantly related parasitoid species

The process of duplication of a gene encoding a protein usually involved in a key physiological process whose function will be hijacked is a major mechanism of toxin recruitment (Fry et al., 2009; Wong & Belov 2012; Casewell et al., 2013). The results of phylogeny and domain analyses indicate that the *Leptopilina* venom RhoGAPs and the calyx RhoGAPs found in *V. canescens* VLPs originated independently from the duplication of the gene encoding the cellular RacGAP1 in these species. The alternative hypothesis of a single initial duplication in the common ancestor of both *Leptopilina* and *V. canescens* would require numerous subsequent loss events since the last common ancestor of both *Leptopilina* and *V. canescens* dates back to the early radiation of parasitoid wasps (Peters et al., 2017). Furthermore, the presence of RhoGAP family proteins in the maternal fluids of parasitoid wasps has not been described outside of these species. This independent convergent recruitment may suggest a similar function in virulence in these two evolutionarily distant parasitoid species. Consistently, our results indicate that venom RhoGAPs in *L. boulardi* and *L. heterotoma* and calyx RhoGAPs in *V. canescens* are associated with extracellular vesicles, named venosomes and VLPs, respectively. While these two types of vesicles have different tissue origins and formation mechanisms, *Leptopilina* venosomes and *V. canescens* VLPs would act as transport systems to deliver virulent proteins, including RhoGAPs, into host hemocytes to alter their function (Pichon et al., 2015; Wan et al., 2019). However, the role of RhoGAPs on host hemocytes is still unclear or even unknown, although we have previously shown that LbmGAP inactivates Rac-like GTPases (Colinet et al., 2007), which are required for successful encapsulation of *Leptopilina* eggs (Williams et al., 2005; Williams et al., 2006).

RacGAP1 is an intracellular multidomain protein consisting of a coiled-coil region followed by a C1 motif (protein kinase C-like zinc finger motif) and a RhoGAP domain (Tcherkezian & Lamane-Vane 2007). *Leptopilina* venom RhoGAPs, on the other hand, possess only a RhoGAP domain preceded by a secretory signal peptide. Sequence comparisons at the genomic level support the hypothesis of a single partial duplication of the RacGAP1 gene in the ancestor of the *Leptopilina* genus, resulting in the loss of a 5’ portion of the sequence encoding the coiled-coil region and the C1 motif, followed by the acquisition of the signal peptide sequence. This initial evolutionary event would then have been followed by further duplication of the ancestral duplicate gene throughout the diversification of the genus *Leptopilina,* leading to the current number of venom RhoGAPs. A first hypothesis for the origin of the signal peptide would be that the loss of part of the 5’ coding sequence in the duplicate resulted in a new N-terminal end for the encoded protein with signal peptide properties. Modification of the N-terminal end of the protein into a fully functional signal peptide may have required some point mutations after partial duplication, as in the case of the antifreeze protein of an Antarctic zoarcid fish (Deng et al., 2010). Another possibility is that the signal peptide originated from a process of partial duplication with recruitment (Katju & Lynch 2006), in which a “proto-peptide signal” coding sequence was acquired from the new genomic environment into which the partial copy was integrated. Finally, it is also possible that the new 5’ region of the duplicated copy results from a chimeric duplication (Katju & Lynch 2006), e.g. from the partial duplication of another gene or by exon shuffling (Vibranovski et al., 2006). The acquisition of the signal peptide probably facilitated the neofunctionalization of the venom RhoGAPs according to the evolutionary model of protein subcellular relocalization (PSR) proposed by Byun McKay & Geeta (2007), according to which a modification of the N-or C-terminal region of a protein can change its subcellular localization, by the loss or acquisition of a specific localization signal, and enable it to acquire a new function. In *V. canescens*, the N-terminal region of the calyx RhoGAPs is more variable in length, with some proteins retaining the C1 motif upstream of the RhoGAP domain and one retaining a coiled-coil region upstream of the C1 motif. Most of the *V. canescens* calyx RhoGAPs were predicted to contain a nuclear localization signal, consistent with (i) the accumulation of VcVLP2 in the nucleus of the calyx cells prior to its association with the virus-derived VLPs (Pichon et al., 2015) and (ii) our results from a proteomic analysis indicating that all 13 calyx RhoGAPs are associated with VLPs. Human RacGAP1 has been described to localize to the cytoplasm and nucleus (Mishima et al., 2002), suggesting that *V. canescens* calyx RhoGAPs have retained the RacGAP1 nuclear localization signal, unlike *Leptopilina* venom RhoGAPs. One of the *V. canescens* calyx RhoGAP genes is located close to the RacGAP1 gene in the genome, suggesting a recent partial tandem duplication, although without support from phylogenetic analyses. The other genes, on the other hand, are scattered throughout the genome suggesting that they originate from one or more older duplication events of the RacGAP1 gene. In contrast to *Leptopilina*, it cannot be excluded that two or more partial duplication events of the RacGAP1-encoding gene occurred in *V. canescens*, followed by further duplication of ancestral duplicates.

### Accelerated evolution through duplication and divergence: pseudogenization and/or neofunctionalization?

Our work revealed a significant divergence for the two multigene families of RhoGAPs in *L. boulardi* and *V. canescens* compared to RacGAP1, illustrated by the presence of substitutions on the arginine essential for GAP activity and/or on one or more of the amino acids shown to be important, or even necessary, for interaction with the Rac GTPases in all RhoGAPs except LbmGAP (and its ortholog LbyGAP) and VcVLP2. The presence of substitutions at key sites in the majority of *L. boulardi* venom and *V. canescens* calyx RhoGAPs suggests a loss of function by pseudogenization. However, peptide matches were found in proteomic analyses for each of these RhoGAPs, indicating that the corresponding genes are successfully transcribed and translated. Furthermore, our results indicate that the ancestrally acquired secretory signal peptide, in which we identified an amino acid under negative selection, is conserved in all *L. boulardi* RhoGAPs and that most *V. canescens* RhoGAPs conserved the nuclear localization signal of RacGAP1. In addition, all RhoGAPs retained the ability to associate with vesicles, namely venosomes in *L. boulardi* and VLPs in *V. canescens*. Finally, we showed that several amino acids embedded in the protein structure evolved under negative selection, suggesting that they are important for protein stability. Taken together, these observations are not consistent with the hypothesis that any of the mutated RhoGAPs is undergoing a pseudogenization process, although it cannot be excluded that this process is quite recent and could have been initiated by the loss of the GAP active site.

An alternative hypothesis to pseudogenization would be functional diversification of the mutated RhoGAPs independent of the Rho pathway. Indeed, we have evidenced an evolution under positive selection for the majority of RhoGAPs in the two multigenic families. Most of the sites under positive selection are located on the surface of the protein and are therefore likely to interact with partners. In addition, directed mutagenesis and two-hybrid experiments in *L. boulardi* have shown that some of these amino acids are not involved in the interaction with Rac GTPases. Although the majority of studies on the RhoGAP domain concern the interaction with Rho GTPases, interactions with other proteins have also been described (Ban et al., 2004; Xu et al., 2013), supporting the hypothesis of a neofunctionalization of the mutated RhoGAPs. Further studies will be needed to determine which functions the mutated RhoGAPs have acquired in relation to parasitism. Recently, a possible role has been proposed for one of the RhoGAPs found in the venom of the ISy line of *L. boulardi* in the induction of reactive oxygen species (ROS) in the central nervous system of *D. melanogaster* in the context of superparasitism avoidance (Chen et al., 2021). However, ROS production is notably regulated by the GTPases Rac1 and Rac2 (Hobbs et al., 2014), whereas this venom protein (named EsGAP1 in Chen et al., 2021 and corresponding to LbyGAP6 in our study) has likely lost its RhoGAP activity due to a substitution on the arginine at position 74. Therefore, the mechanism by which EsGAP1 would be involved in ROS induction is unclear.

Other potential factors involved in parasitic success have been described previously that are encoded by large gene families and similarly correspond to truncated forms of cellular proteins that have retained only a single conserved domain, such as protein tyrosine phosphatases (PTPs) or viral ankyrins (V-ANKs) of bracoviruses. In the case of bracovirus PTPs, some are active as phosphatases, while others are mutated in their catalytic site and have been shown to be inactive as PTPs (Provost et al., 2004). In the latter case, as well as for V-ANK, it has been suggested that by binding to their target, they may act as constitutive inhibitors of the function of the corresponding cellular protein (Provost et al., 2004; Thoetkiattikul et al., 2005). Moreover, the model of adaptive evolution by competitive evolution of duplicated gene copies (Francino 2005) predicts that after a first step in which different copies explore the mutation space, once a protein with an optimal function is obtained, the other copies will begin to decay and undergo pseudogenization. This can lead to intermediate situations where both functional and non-functional proteins are produced, as observed for different virulence protein families of parasitoid wasps. Some of the described features of *L. boulardi* and *V. canescens* RhoGAPs may suggest that a process of competitive evolution is underway in *V. canescens* and *L. boulardi*, although our results are also consistent with the hypothesis of a possible neofunctionalization of these vesicle-associated proteins as discussed above.

In conclusion, we evidenced the independent convergent origin and accelerated evolution of a multigene family of vesicle-associated RhoGAP proteins in two unrelated parasitic wasps. Strikingly, these vesicles, which are similarly involved in parasitism, are produced in distinct organs: the venom apparatus in *Leptopilina* and the ovarian calyx in *V. canescens*. In the case of *Leptopilina* RhoGAPs, the acquisition of a secretory signal peptide after incomplete duplication of the RacGAP1 gene allowed secretion into the venom where the vesicles are formed. *V. canescens* RhoGAPs, on the other hand, probably retained the RacGAP1 nuclear localization signal, because it is in the nucleus of calyx cells that VLPs are formed. Another striking point is that our results suggest a possible functional diversification of vesicle-associated RhoGAPs in both species, with the exception of LbmGAP (and its ortholog LbyGAP) in *L. boulardi* and VcVLP2 in *V. canescens*, which are probably the only ones with RhoGAP activity. An open question would be whether all RhoGAPs are important for parasite success, but by different mechanisms, independent of the Rho pathway or not, depending on whether RhoGAPs are mutated or not.

## Acknowledgements

We are highly grateful to Christian Rebuf for help in insects rearing. We also thank Pr. M. T. Kimura (Hokkaido University, Japan) for providing biological material. We would also like to thank Anthony Bretaudeau (INRIA, Rennes, France) for help in the use of the workflows available at the BIPAA platform, the genotoul bioinformatics platform Toulouse Occitanie (Bioinfo Genotoul, https://doi.org/10.15454/1.5572369328961167E12) for providing computing resources and the BIG bioinformatics platform from the PlantBios infrastructure for providing facilities and technical support. Preprint version 3 of this article has been peer-reviewed and recommended by Peer Community In Genomics (https://doi.org/10.24072/pci.genomics.100248; Bravo, 2021).

## Data and supplementary information availability

Raw data from the Illumina RNASeq sequencing of the *L. victoriae* venom apparatus are available at NCBI under the BioProject ID PRJNA974978.

The mass spectrometry proteomics data from *L. boulardi* and *L. heterotoma* venosomes have been deposited to the ProteomeXchange Consortium via the PRIDE (Perez-Riverol et al., 2022) partner repository with the dataset identifier PXD041695.

Supplementary information and material are available online at Data INRAE: https://doi.org/10.57745/K82IWQ

## Conflict of interest disclosure

The authors declare that they comply with the PCI rule of having no financial conflicts of interest in relation to the content of the article.

Jean-Luc Gatti and Marylène Poirié are recommenders of PCI Zoology.

